# Copepod life history evolution under high and low food regimes

**DOI:** 10.1101/2022.12.09.519570

**Authors:** Alexander Blake, Dustin J. Marshall

## Abstract

Copepods play a critical role in the carbon cycle of the planet – they mediate the sequestration of carbon into the deep ocean, and are the trophic link between phytoplankton and marine foodwebs. Global change stressors that decrease copepod productivity create the potential for catastrophic positive feedback loops. Accordingly, a growing list of studies examine the evolutionary capacity of copepods to adapt to the two primary stressors associated with global change: warmer temperatures and lower pH. But the evolutionary capacity of copepods to adapt to changing food regimes, the third major stressor associated with global change, remains unknown. We used experimental evolution to explore how a 10-fold difference in food availability affects life history evolution in the copepod, *Tisbe sp*. over two years, and spanning 30+ generations. Different food regimes evoked evolutionary responses across the entire copepod life history: we observed evolution in body size, size-fecundity relationships and offspring investment strategies. Our results suggest that changes to food regimes reshape life histories and that cryptic evolution in traits such as body size is likely. We demonstrate that evolution in response to changes in ocean productivity will alter consumer life histories, and may distort trophic links in marine foodchains. Evolution in response to changing phytoplankton productivity may alter the efficacy of the global carbon pump in ways that have not been anticipated until now.1

## Introduction

The oceans account for around 50% of carbon sequestration (Sabine *et al*. 2004). Phytoplankton are the primary producers that account for most marine carbon fixation (del Giorgio & Duarte 2002), but only a small proportion is sequestered into the deep sea (Dunne *et al*. 2007). Copepods play a key role here, they feed on phytoplankton, and excrete them in faecal pellets that sink much faster than individual phytoplankton cells. Of the total biomass fixed every year, up to 15-58% is consumed by copepods during some times of the year, and 7.4-29 gigatonnes of carbon are annually consumed on average (Steinberg & Landry 2017). Given the massive biomass turnover between phytoplankton and copepods, anything that affects the dynamics of this trophic link has significant consequences for the efficacy global carbon pump. If global change affects copepods, then the carbon pump will also be affected. Worse still, a positive feedback loop could be generated: increasing atmospheric CO_2_ increases stressors (e.g. higher temperatures, lower seawater pH), decreasing copepod productivity, decreasing carbon sequestration rates, leading to more rapid increases in atmospheric CO_2_. The capacity for copepods to adapt to global change represents a key uncertainty in making predictions for the future of the global carbon pump.

For organisms with relatively short lifespans, predicting their capacity to adapt to global change requires an understanding of their evolutionary capacity to adapt to future conditions (Munday *et al*. 2013; Kelly & Griffiths 2021). Accordingly, a growing number of studies estimate how copepods adapt to higher temperature regimes and lower pH – the two primary stressors associated with global change in the sea (Brennan *et al*. 2021, 2022; deMayo *et al*. 2021; Sasaki & Dam 2021). While temperature and pH are undoubtedly important, they are not the only factors that will change in future oceans – food availability is likely to be dramatically different for copepods under future regimes (Fu *et al*. 2016), and indeed is already changing (Capuzzo *et al*. 2018).

Phytoplankton depend on sunlight for photosynthesis, so they are most productive in the surface waters of the ocean – the productivity of these surface waters are likely to alter with global change. The upper, sunlit layers of the ocean are depleted of nutrients by growing phytoplankton and must replenished by cooler, nutrient-rich water from deeper, darker layers. Unfortunately, global warming can create more stratification in the water column, generating more intense thermoclines, and reducing the rate of mixing between water layers (Hannon *et al*. 2001). This thermal shoaling is likely to reduce phytoplankton productivity in some places by up to 40% (Jang *et al*. 2011). On the other hand, some regions are predicted to experience more intense storms, which will actually enhance mixing of surface waters with deeper layers, possibly increasing the productivity in those places (Nicholson *et al*. 2016). Thus, copepods must not only evolve to cope with warmer, more acidic oceans, they are also likely to experience massive changes to their food regimes. The capacity for evolution in response to such changes remains unclear.

Food regimes have classically been recognised as key drivers of life history evolution across a broad range of species (Macarthur & Levins 1967; Tilman 1982; Chesson 2000; Grant & Grant 2006), but the direction and nature of such effects remain debated. For example, guppies in food-scarce environments evolve larger body sizes (Felmy et al. 2022) while flies get smaller (Kolss et al. 2009). Furthermore, in some cases the direction of evolution is opposed to the direction of plasticity imposed by the environment (Potter et al. 2021). Because this countergradient variation (*sensu* Conover & Schultz 1995) obscures evolutionary responses in field measurements, we know little about how widespread it might be in other natural systems, particularly in response to food (Conover et al. 2009). As such, it is difficult to predict how, or even if, copepods will evolve in response to changes in food based on studies in other systems. Biogeographical studies are similarly ambiguous. For example, some imply that copepods should become larger under low productivity regimes but whether body size is decreasing because of food availability *per se* or other covarying factors such as predation risk remains unclear (Brun et al. 2016). As far as we are aware, no study has directly addressed the capacity of copepods to evolve in response to different food regimes. We sought to address this important knowledge gap here.

We subjected copepod populations to either high-food or low-food environments for 2 years and then used multigenerational common garden experiments to examine how life histories have evolved independently of any cross-generational parental effects (Burgess & Marshall 2014) and detect countergradient variation (Conover & Schultz 1995). We evaluated key life history traits that have responded to food regime in other species (e.g. Trinidad guppies: Felmy et al. 2022), including size and age at maturity, egg size, and fecundity, accounting for the effects of maternal size on reproductive traits due to potential correlations between egg size, fecundity, and mother size (Moran & McAlister 2009; Barneche et al. 2018). We did so with a novel study species from the genus *Tisbe*, a group of copepods that is highly suited to long-term experimental evolution due to a. its hardiness and well-documented husbandry (Webb & Marcotte 1984; Arndt & Sommer 2014); b. its geographic ubiquitousness and therefore suitability for follow-up studies along natural primary productivity gradients (*GBIF Secretariat* 2022); and c. exposure to primary productivity that can vary by up to an order of magnitude in its natural range (Beardall et al. 1997).

## Materials and Methods

### STUDY ORGANISM

*Tisbe sp*. is a littoral marine copepod from the Tisbidae family (Arthropoda: Harpacticoida) that has not been resolved to species level in the Southern Ocean. Mothers brood clutches of approximately 10-40 eggs on abdominal egg-sacs for 1-3 days. The naupliar larvae are facultative planktotrophs and pass through 6 stages before metamorphosing into a juvenile copepodid approximately 3-5 days after hatching with *ad libitum* food. Juveniles pass through a further 6 copepodid stages before reaching the final adult and sexually mature stage at approximately 18-20 days old. We collected approximately 5000 copepods from Brighton Marina in Port Phillip Bay, Australia in May 2017. We isolated and maintained these ‘ancestral’ populations in gently oxygenated 500 ml mason jars in freshly pasteurised seawater (FSW). Ancestral populations were reared on the marine microalga *Dunaliella tertiolecta* and fed at a rate of 2.475 × 10^9^ algae cells per litre of copepod culture per week.

*Dunaliella* cultures were maintained using F2 media (Guillard & Ryther 1962) and algae concentrate for feeding copepod populations was prepared three times a week. Density of algal cultures was determined spectroscopically using a SPECTROstar Nano and algal concentrations were adjusted to approximately 1.1 × 10^10^ *D. tertiolecta* cells/L in FSW after centrifuging and removing media. Ancestral populations were fed manually 3 times a week, while for experimental cultures housed in flow-through culture vessels feeding was automated (see Experimental Evolution). See Figure S2 of a schematic overview of the series of experiments we conducted during the study.

### EXPERIMENTAL EVOLUTION

#### Experimental design

Experimental evolution commenced on the 13^th^ of October 2018 at Monash University Clayton Campus, Melbourne, Australia. Copepods from ancestral stocks were randomly assigned to either high-food or low-food environments, which differed in their rate of food supply (of *D. tertiolecta cells*) by an order of magnitude. Each population was founded with approximately 1000 individuals using a Folsom plankton splitter to ensure copepods were randomly assigned, and the 1L glass culture vessels were topped up with FSW. In total, 20 copepod cultures were subjected to experimental evolution, consisting of 10 high-food and 10 low-food replicates reared in separate glass pressure-equalising dropping funnels. One low-food replicate went extinct one year into the experiment due to bacterial contamination.

High and low food rates were determined through pilot experiments in 2018 and were set as follows: high food replicates received 4.5 × 10^9^ algae cells per litre of copepod culture per week, and low food replicates received 4.5 × 10^8^ algae cells per litre of copepod culture per week. An intermediate feeding rate (2.475 × 10^9^ algae cells per litre of copepod culture) was used for the ancestral populations and the common garden. Differential food supply in experimental treatments was ramped up gradually, with high and low food treatments receiving the same intermediate rate of supply for the first week, partial treatments in week two (3.5 × 10^9^ and 1.5 × 10^9^ *D. tertiolecta* cells per week, respectively), and final treatments of high or low food supply by week three after initiating the experiment. These treatment differences were then maintained for the next 16 months.

Experimental cultures were organised into blocks of four cultures (in a randomised sequence of 2 high-food and 2 low-food replicates) due to spatial constraints, with five such blocks in total. Each block received food from a separate algae reservoir using a Kamoer X4 peristaltic dosing pump. Both high-food and low-food treatments received a total inflow of 80mL per litre of culture per day (on weekdays only, no dosing on weekends). High-food treatments were dosed with 80mL of algae concentrate per day, while low-food treatments received 8 mL of algae concentrate and 72mL of FSW (i.e. 10% of the high food regime). Pumps provisioned algae concentrate semi-continuously by dosing 12 times a day at 2-hour fixed intervals, and were recalibrated every 2 months. Pilot work indicated that very few adult copepods were lost at this rate of inflow-outflow. Algae and FSW reservoirs were covered to minimise evaporation and cleaned weekly to minimise build-up of waste and pathogens, and algae was kept well-mixed using simple magnetic stirrers at low speed. Laboratory temperature was set at 21°C with a light:dark photoperiod of 12h:12h, and salinity was maintained at 37 ppt with monthly monitoring.

### COMMON GARDENS

#### Experimental design

To disentangle genetic responses from plastic responses in experimental evolution, individuals need to be sampled from divergent evolutionary lineages and reared in a common environment (Huey & Rosenzweig 2009). Because environmental effects can persist between generations, such common environment (or ‘common garden’) experiments must also be performed over multiple generations to minimise any lingering parental and grandparental effects on offspring phenotypes (Burgess & Marshall 2014). To evaluate the evolutionary response to high- and low-food environments, we performed a common garden experiment wherein copepods were sampled from their treatment cultures (G0) and their descendants were reared (separately) under the same environmental conditions over two generations (G1 and G2).

Paired high- and low-food cultures were randomly sampled between the 18^th^ and 20^th^ of February 2020. 10 gravid G0 mothers were collected from each culture, photographed, and transferred to sterile plastic culture trays containing 4 mL FSW. 16uL of 10000 units mL^-1^ (approximately 6 mg mL^-1^) penicillin G and 10 mg ml^-1^ streptomycin solution (Sigma-Aldrich) was added to each tray to inhibit the growth of pathogens (Gangur & Marshall 2020). G0 mothers were monitored daily and returned to their cultures after releasing their G1 eggs, until all eggs had hatched and all G0 mothers had been removed (generally 3-5 days after initial collection). All common garden replicates commenced G1 with >100 larvae.

Throughout the experiment, *D. tertiolecta* was provisioned at an intermediate level of food supply. For G0, G1, and G2 juveniles and adults, food was provisioned each weekday (5 times a week) in a 176 uL pulse from a 1.1 × 10^7^ cells/mL stock representing an intermediate food supply of approximately 2.475 × 10^9^ cells per litre per week. Due to their lower feeding rate, larvae were provisioned a single 176uL (at 1.1 × 10^7^ cells/mL) pulse of food to achieve the same maximum ambient food density experienced by adults under an intermediate feed regime (approx. 5 × 10^6^ *D tertiolecta* cells per mL each day).

With a generation time of ∼17 days, our *Tisbe sp*. cultures had undergone approximately 30 generations of evolution prior to common gardening, which commenced on 18 February 2020. Evolutionary responses in five traits were measured across the three generations.

#### Data collection

Maternal size, mean egg size, and fecundity were measured in G0, G1, and G2, while survival and age at maturity were measured in G1 and G2 only. Maternal size was measured as length between end of urosome to tip of prosome. We used mean egg size from 10 randomly measured eggs within each clutch, and also estimated fecundity as number of eggs per egg sac. Maternal body size, egg size, and fecundity were recorded with photographs using a Motic Moticam 1080 camera mounted on an Olympus SZ61 dissecting microscope and digitally measured using FIJI version 1.53c (Schindelin et al. 2012).

Freshly hatched G1 larvae within each replicate were counted and randomly allocated to individual culture trays of 25 ± 5 individuals with 4 mL FSW, antibiotics, and food. When metamorphosis was first observed within a replicate, the replicate was censused and individuals were transferred to fresh trays with 4 mL FSW, antibiotics, and food. For juveniles, water was then changed and censusing was conducted weekly until sexual maturity was first observed within a replicate. Then subreplicates were censused, pooled, mixed, and reallocated into new adult subreplicates of 25 ± 5 with fortnightly water changes and censusing. All reproductive mothers were collected and photographed with their first clutch. Mothers and their egg sacs were photographed, then pooled at the replicate level into fresh culture trays containing 4 mL FSW, antibiotics, and food. At least 5 G1 mothers and 50 offspring were obtained for 17 of 19 replicates (8 high-food and 9 low-food cultures), and these G2 offspring were collected for the final stage of the experiment. Replicates containing copepods from the two remaining high-food cultures were accidentally dropped before reaching G2.

G2 larvae were collected from culture trays containing gravid G1 mothers in a similar fashion to G1 larvae collection from G0 mothers, but we also accounted for temporal staggering. For each replicate, G2 larval collection took place over a week after the first clutch hatched. Larvae collected during this week were continually transferred to culture trays at a density of 25 ± 5 individuals with 4 mL FSW, antibiotics, and food. At the end of the collection week these larvae (some of which had metamorphosed) were re-pooled, mixed, and randomly allocated to new culture trays of 25 ± 5 individuals in 4 mL FSW, antibiotics, and food. G2 larvae that hatched outside this initial collection week were retained but reared in separate trays. G2 larvae were then reared to sexual maturity and reproductive G2 mothers were measured following the same protocol used for G1.

### STATISTICAL ANALYSES

All phenotypes were analysed using linear mixed effects models. Full models for size, survival, and age at maturity included treatment and generation as fixed effects, as well as their interaction. Fecundity and egg size were modelled separately for each generation due to complex interactions and included mother size as a fixed covariate, as well as its interaction with treatment. Using model predictions for egg size and fecundity, reproductive volume was calculated for G2 *a posteriori* as:

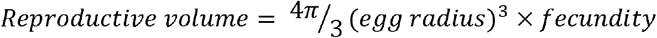

G0 feeding block was included as a fixed effect in all models due to insufficient replication to treat as a random effect. All models also included culture nested within treatment as a random intercept term. Where interaction terms were nonsignificant they were removed and the analysis was repeated. Models were evaluated using type III tests due to imbalance of high-food and low-food replicates. *p* values for relevant fixed effects were obtained with *F* tests using Sattertwaithe’s approximation (Kuznetsova *et al*. 2017).

Analyses were performed with R version 4.1.2 (R Core Team 2021) and RStudio version 2021.09.1 (RStudio Team 2021), using dplyr (Wickham et al. 2021) to prepare the data. Linear mixed effect models were fitted with lme4 (Bates et al. 2015). The lmerTest package (Kuznetsova et al. 2017) was used to perform Sattertwaithe’s approximations and type III tests on fixed effects, and likelihood ratio tests on random effects. Bootstrapped 95% confidence intervals were obtained between cultures using merTools (Knowles & Frederick 2020), and plots were built using ggplot2 (Wickham 2016). Diagnostic residuals vs fits and QQ plots were visually assessed as per Keough & Quinn (2002), and VIF calculated to check for collinearity. Predictors were plotted against each other to visually assess acceptable domain/range overlap.

## Results

### PHENOTYPES OBSERVED PRIOR TO COMMON GARDENING (G0)

Copepods reared in high- and low-food environments (G0) differed slightly in body size (Figure 1a) and fecundity (Figure 1d). Copepods reared with high food provisioning were slightly larger than those reared with low food provisioning, shown by the significant interaction between generation and food lineage (Table 1) and cell means (Figure 1a). Consequently, copepods with high food provisioning also tended to have slightly higher fecundity due to the significant positive covariance between fecundity and maternal size (Table 4), although there was no main effect of food lineage on fecundity.

**Table 1:** Linear mixed effects model for the effect of evolutionary treatment on adult female body size across three generations (G0, G1, G2), with ancestral G0 cultures reared within feeding blocks of four. *p* values are provided for tests of interest, significant effects are specified in bold, and random effect is specified in italics. Degrees of freedom (df) reported as numerator df, denominator df. n_low_= 9 and n_high_ = 10.

| Effect | Df | Mean squares | F | *p* |
| --- | --- | --- | --- | --- |
| Treatment | 1, 11.67 | $5.69 \times 10^{-1}$ | 0.16 | 0.70 |
| Generation | 2, 691.43 | $3.20 \times 10^{-4}$ | 287.52 | **<10<sup>-15</sup>** |
| Block | 4, 11.76 | $1.95 \times 10^{-3}$ | 0.98 | 0.45 |
| Treat x Gen | 2, 686.54 | $1.01 \times 10^{-2}$ | 5.10 | **0.006** |
| *Culture(Treat)* | | $1.93 \times 10^{-3}$ | | |

**Figure 1:**
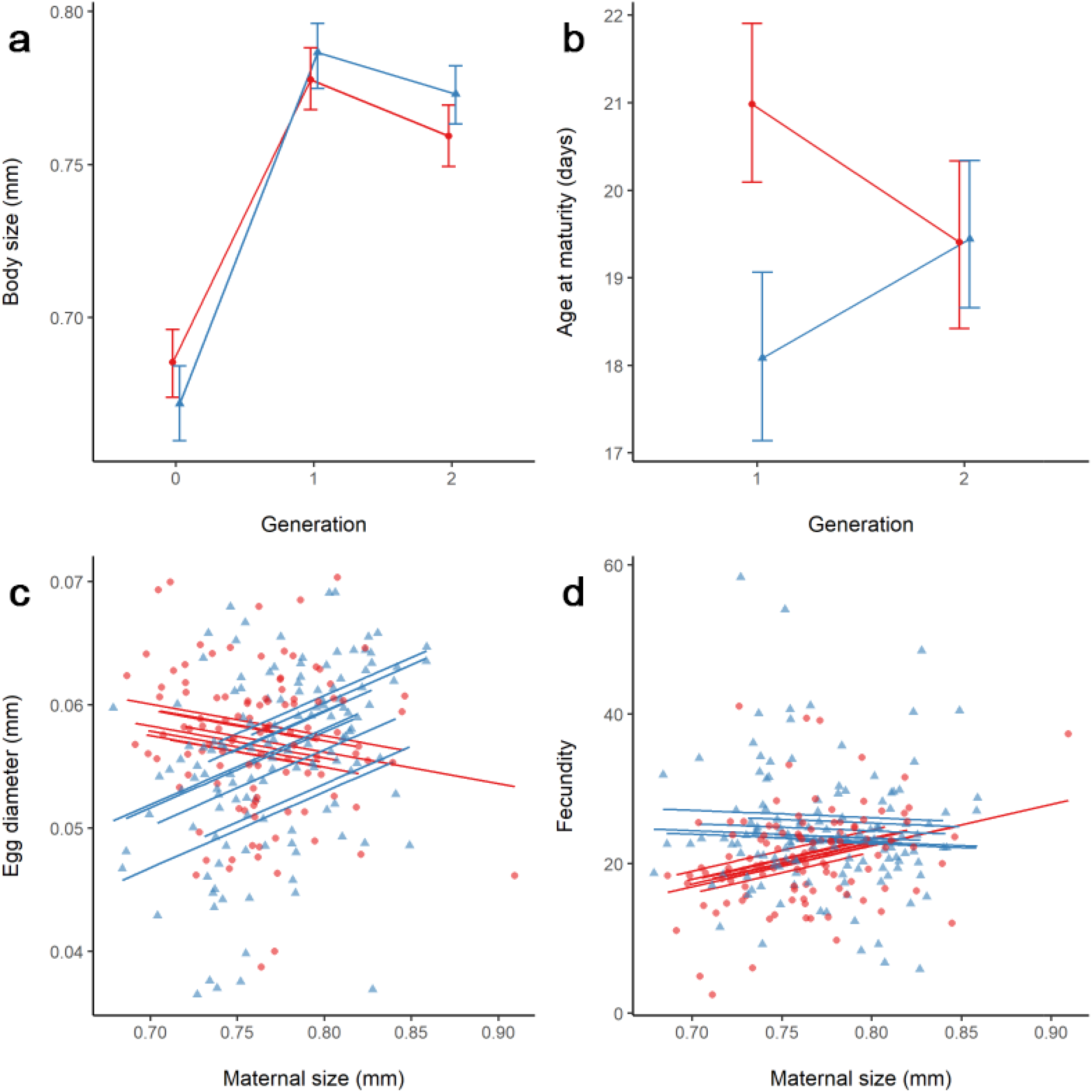
Key life history responses to high (red, circular points) and low (blue, triangular points) food regimes. Panels show a: mean adult female body length across all three generations (n_low_= 9, n_high_= 10), b: mean cohort age at first observation of sexual maturity (within each replicate) in G1 and G2 (n_low_ = 9, n_high_ = 9), c: mean first-clutch egg diameter in G2 mothers (n_low_ = 9, n_high_ = 8), d: mean first-clutch fecundity in G2 mothers (n_low_ = 9, n_high_ = 8). Errors bars show between-culture bootstrapped 95% confidence intervals (a, b). Points show raw data at the sub-replicate (individual female) level, lines show regressions at the culture level (c, d).

### SURVIVAL AND BODY SIZE

The significant interaction between generation and food lineage indicates that the effect of food lineage changed across generations (Table 1). In the common-gardened generations (G1 and G2), females were larger in the low-food regimes relative to the high-food regimes (Figure 1a). There was also an increase in body size over the course of the common gardening relative to G0. Survival in the common garden was unaffected by food lineage, showing no significant effect (Table S1 and Figure S1, see Supporting Information).

### AGE AT MATURITY

Age at maturity differed between food regimes in G1 copepods but converged in G2, shown by the significant interaction between food lineage and generation (Table 2). Plotted means indicate that in G1, copepods from high-food lineages matured later despite being smaller than copepods from low-food lineages, but by G2 these differences had disappeared (Figure 1b).

**Table 2:** Linear mixed effects model for the effect of evolutionary treatment on age at maturity across two generations (G1, G2), with ancestral G0 cultures reared within feeding blocks of four. *p* values are provided for tests of interest, significant effects are specified in bold, and random effect is specified in italics. Degrees of freedom (df) reported as numerator df, denominator df. n_low_= 9 and n_high_ = 9.

| Effect | Df | Mean squares | F | *p* |
| --- | --- | --- | --- | --- |
| Treatment | 1, 10.26 | 95.65 | 6.31 | **0.03** |
| Generation | 1, 517.99 | 2.39 | 0.16 | 0.69 |
| Block | 4, 10.44 | 15.20 | 1.00 | 0.45 |
| Treat x Gen | 1, 517.91 | 517.91 | 18.33 | **<10<sup>-4</sup>** |
| *Culture(Treat)* |  | 14.74 |  |  |

### REPRODUCTIVE OUTPUT

The relationships between maternal size and reproduction evolved in response to food regime, but the effect of food lineage was only significant in the second generation of common gardening (Egg size: Table 3; Fecundity: Table 4). In G2, larger mothers from the low-food lineages produced larger (Figure 1c) but slightly fewer offspring (Figure 1d), whereas from the high-food lineages larger mothers produced smaller (Figure 1c) but many more offspring (Figure 1d). Combining these two components of reproduction (Figure 2), mothers from the low-food lineages exhibited a steeper positive relationship between body size and reproductive volume than mothers from the high food lineages (with slopes of 6.64 × 10^−2^ vs 3.51 × 10^−2^ respectively).

**Table 3:** Linear mixed effects models for the effects of evolutionary treatment and maternal size on mean egg size across three generations (G0, G1, G2), with ancestral G0 cultures reared within feeding blocks of four. *p* values are provided for tests of interest, significant effects are specified in bold, and random effect is specified in italics. Degrees of freedom (df) reported as numerator df, denominator df. n_low_= 9 in G0-2 and n_high_ = 10, 9, and 8 in G0, G1, and G2 respectively.

| Effect | Df | Mean squares | F | *p* |
| --- | --- | --- | --- | --- |
| **G0** |  |  |  |  |
| Treatment | 1, 12.98 | $1.36 \times 10^{-4}$ | 3.00 | 0.11 |
| Maternal Size | 1, 77.59 | $7.36 \times 10^{-5}$ | 1.62 | 0.21 |
| Treat x Size | 1, 95.48 | $3.12 \times 10^{-5}$ | 0.69 | 0.41 |
| Block | 4, 12.45 | $3.87 \times 10^{-5}$ | 0.85 | 0.52 |
| *Culture(Treat)* | | $4.14 \times 10^{-5}$ | | |
| **G1** |  |  |  |  |
| Treatment | 1, 12.32 | $1.83 \times 10^{-6}$ | 0.07 | 0.80 |
| Maternal Size | 1, 184.00 | $2.24 \times 10^{-5}$ | 0.84 | 0.36 |
| Treat x Size | 1, 184.00 | $1.39 \times 10^{-5}$ | 0.52 | 0.47 |
| Block | 4, 13.11 | $8.57 \times 10^{-6}$ | 0.32 | 0.86 |
| *Culture(Treat)* | | $2.48 \times 10^{-5}$ | | |
| **G2** |  |  |  |  |
| Treatment | 1, 232.57 | $6.11 \times 10^{-4}$ | 17.69 | **<10<sup>-4</sup>** |
| Maternal Size | 1, 231.65 | $9.87 \times 10^{-4}$ | 2.85 | 0.09 |
| Treat x Size | 1, 233.32 | $5.99 \times 10^{-3}$ | 17.32 | **<10<sup>-4</sup>** |
| Block | 4, 11.35 | $9.57 \times 10^{-4}$ | 2.77 | 0.08 |
| *Culture(Treat)* | | $3.25 \times 10^{-5}$ | | |

**Table 4:** Linear mixed effects models for the effects of evolutionary treatment and maternal size on fecundity across three generations (G0, G1, G2), with ancestral G0 cultures reared within feeding blocks of four. *p* values are provided for tests of interest, significant effects are specified in bold, and random effect is specified in italics. Degrees of freedom (df) reported as numerator df, denominator df. n_low_= 9 in G0-2 and n_high_ = 10, 9, and 8 in G0, G1, and G2 respectively.

| Effect | Df | Mean squares | F | *p* |
| --- | --- | --- | --- | --- |
| **G0** |  |  |  |  |
| Treatment | 1, 11.15 | 0.83 | 0.05 | 0.83 |
| Maternal Size | 1, 102.98 | 206.72 | 11.38 | **0.001** |
| Treat x Size | 1, 114.55 | 3.11 | 0.17 | 0.68 |
| Block | 4, 10.98 | 10.98 | 0.48 | 0.75 |
| *Culture(Treat)* |  | 16.10 |  |  |
| **G1** |  |  |  |  |
| Treatment | 1, 9.99 | 8.75 | 0.27 | 0.62 |
| Maternal Size | 1, 183.48 | 863.73 | 26.38 | **<10<sup>-6</sup>** |
| Treat x Size | 1, 183.95 | 0.99 | 0.03 | 0.86 |
| Block | 4, 10.57 | 9.39 | 0.29 | 0.88 |
| *Culture(Treat)* |  | 30.30 |  |  |
| **G2** |  |  |  |  |
| Treatment | 1, 234 | 383.72 | 7.44 | **0.007** |
| Maternal Size | 1, 234 | 165.44 | 3.86 | 0.051 |
| Treat x Size | 1, 234 | 342.10 | 6.69 | **0.01** |
| Block | 4, 234 | 271.44 | 5.36 | **<10<sup>-3</sup>** |
| *Culture(Treat)* |  | 49.71 |  |  |

**Figure 2:**
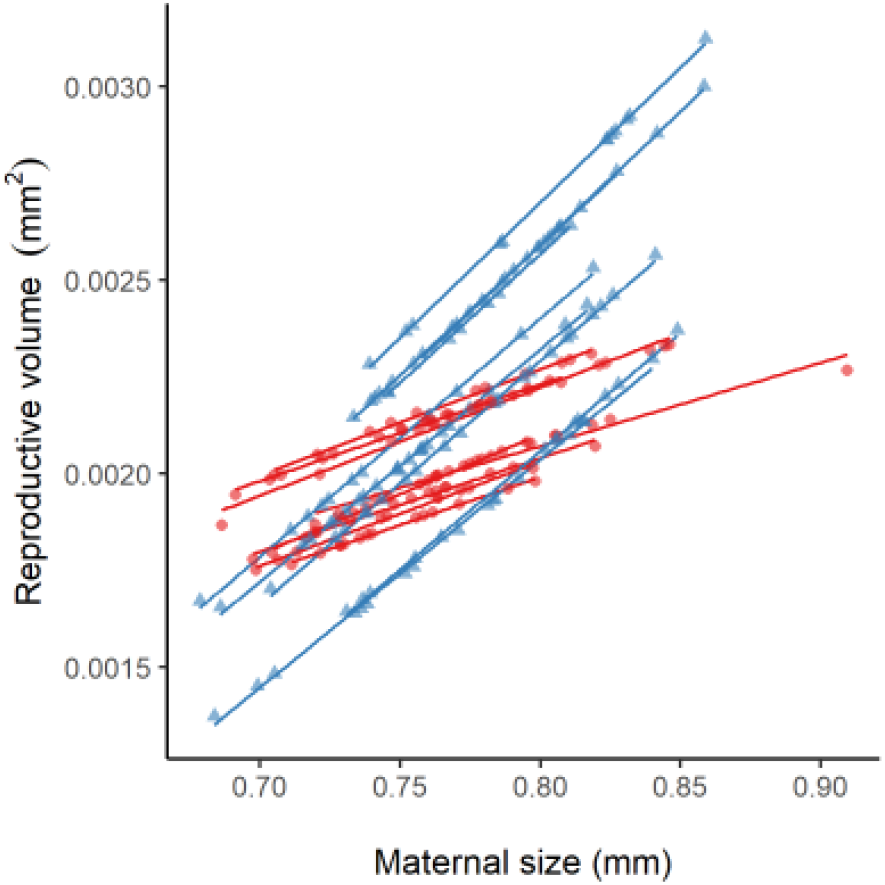
Predicted relationships between reproductive volume (product of mean egg volume and fecundity) and maternal size in G2 mothers from high-food (solid red lines, circular points) and low-food (dotted blue lines, triangular points) regimes. Points show data at the individual level, lines show regressions at the culture level (n_low_=9,n_high_=8), data obtained from G2 egg-size and G2 fecundity model coefficients.

## Discussion

Copepods under different resource regimes evolved different life histories: body size, fecundity, and per-offspring investment all evolved, while age at maturity also changed but appeared to be driven by a strong parental environment effects that dissipated across generations. Copepods evolved to be slightly larger in low food lineages, and within those lineages larger mothers invested more in their individual offspring. In high food lineages, copepods evolved to be smaller and within those regimes larger mothers invested less in their individual offspring but were much more fecund. Interestingly, we found evidence for differences in evolved responses to different resource environments relative to the expressed phenotypes in those environments, indicating countergradient evolution. Overall, our results suggest that the changes in food regimes predicted to occur in future oceans will generate life history evolution in copepods but not in straightforward ways. Our results also imply that biogeographical patterns in life history and covariance between productivity and phenotypes may provide very little information or predictive power about the underlying genetic clines in these traits due to countergradient variation.

Copepods were smaller when reared in a low-food environment, but their offspring grew to be larger when transferred to an intermediate-food common environment, indicating countergradient evolution in body size. Assuming cubic scaling with length, copepods from low-food environments were only 5.9% smaller by volume. Once released from the low-food conditions, they were 5.6% larger than copepods from high-food lineages, suggesting that the impact of food scarcity was moderated by genetic compensation (Grether 2005). Such countergradient variation is observed in field studies as well and seems particularly common in fish (Arendt & Wilson 1999; Conover et al. 2009). We suggest that global changes to phytoplankton productivity will evoke evolutionary change in copepod body sizes but that these changes may be masked by countergradient evolution. Studies seeking to understand how copepod body sizes have changed, and continue to change should consider common garden experiments to disentangle phenotypic and genetic responses, which may counteract each other, resulting in what is sometimes called ‘cryptic evolution’ (Grether 2005).

The relationship between body size and reproductive investment evolved, albeit in subtle ways. In low food environments, egg size increased with maternal size at the expense of fecundity while, while clutch size increased with maternal size at the expense of egg size in high-food lineages. These results are in keeping with general offspring size theory whereby in poor environments, mothers make larger offspring in order to buffer them from harsh conditions (Parker & Begon 1986). Such phenotypic effects have been observed in other taxa (e.g. Allen et al. 2008; Fox & Czesak 2000; see Marshall et al. 2018 for a review) but to our knowledge, ours is one of the few unequivocal demonstrations that differences in reproductive investment strategies can rapidly evolve.

The way reproduction scaled with maternal size showed clear evolutionary responses to food regime. Regardless of evolutionary lineage, larger mothers reproduced more than smaller mothers, implying larger mothers have more resources to invest in offspring (whether nutrients or energy is unclear, but an interesting question for future studies). Larger individuals may be better competitors, able to acquire more resources for reproduction (e.g. Bassar et al. 2016). Larger mothers may also alter their allocation of resources among fitness components, or may simply be physically able to brood more or larger eggs (Bernardo 1996). We find that the way in which larger mothers deploy this resource advantage relative to smaller mother depends on the resource regime in which they evolved – in low-food regimes, they make better provisioned offspring, in high-food regimes, they make more offspring. These different allocation patterns yield an evolved difference in reproductive scaling – we found *a posteriori* that reproductive scaling was steeper in low resource environments than high resource environments. Interestingly, low resource lineages also evolved larger body sizes (at least genetically) – it may be that they evolved to be larger so as to gain the fitness advantages of increased body size that come from steeper reproductive scaling, but this remains speculative. Nevertheless, to our knowledge our study is the first to demonstrate experimental evolution in reproductive scaling, which has clear consequences for population dynamics (Marshall et al. 2022) and should be a focus in future work. Generally theory predicts that when reproductive scaling is steeper (i.e. the slope of reproductive volume or output against maternal size is steeper), population replenishment and productivity depends more strongly on having larger females in population (Marshall et al. 2022). Given smaller body sizes are expected under higher temperatures (Atkinson & Sibly 1997), our results suggest that combining high temperature regimes (the redaction of larger females) and low food conditions (steeper reproductive scaling) could synergise to generate catastrophic reductions in population productivity. An important next step will be to examine how combined stressor regimes (e.g. temperature and food) affect evolutionary trajectories.

We found differences in the timing of maturity in G1 copepods but not G2, suggesting that age at maturity is a transgenerational plasticity effect rather than an evolved response. It seems to us at least that the low food parental environment programs offspring to mature sooner than offspring whose parents experience high food levels. While transgenerational plasticity in key life history traits is relatively common (Yin *et al*. 2019), we are unaware of studies that have shown such effects on the timing of maturity specifically. That this effect disappears after a single generation suggests that this parentally program trait has evolved to track environmental variation closely, with minimal persistent lags as predicted by some theory (Burgess & Marshall 2014).

Our study faced several noteworthy limitations due to the intensity of work involved. First, we only use two generations for common gardening, testing for parental and grandparental effects. While more distant ancestors may also influence phenotypes, such experiments become increasingly difficult and indeed are rare in the literature (Yin *et al*. 2019). Drawing the line at grandparental effects was a necessary compromise which was arbitrary but generally considered sufficient for experimental evolution studies (Garland & Rose 2009). Further, successive generations of common gardening can impose a new selection pressure diluting the original evolutionary signal of interest. Second, we had limited logistical capacity to maintain experimental replicates. Consequently, we decided to maximise replication in the food treatments by omitting an intermediate-food ‘control’ line. Nor did we control for the use of antibiotics, which was considered necessary to minimise pathogenic bacteria and maximise copepod sample size. Though penicillin G and streptomycin in low doses have shown minimal deleterious impact on *Tisbe* (Gangur & Marshall 2020) and other arthropods (e.g. *Drosophila*: Heys *et al*. 2018), copepod gut flora may have been affected with knock-on impacts on life history traits. Overall, we were limited to testing the relative divergence of life histories in high- and low-food lineages, assuming no interactive effects of antibiotics. Third, we were similarly limited in our ability to control all environmental parameters. Notably, we did not attempt to control pH or population densities, although informal monitoring of the latter suggested that high- and low-food populations rapidly reached equilibria around 20000 and 5000 individuals respectively and oscillated around these carrying capacities for the remainder of the experiment. In general we did not attempt to discern exact drivers of life history changes in the present study, and given that food provisioning in the evolutionary lines differed by an order of magnitude but population densities did not, we suspect that population density likely plays a role. Identifying the precise mechanisms driving evolution in high- and low-food regimes should be a major priority for future work.

In summary, we show that differing food regimes induce rapid evolutionary responses relative to rate and magnitude of anthropogenic change that may induce those responses (Tagliabue et al. 2021), affecting every aspect of their life history from offspring size, through to growth and reproduction. These evolutionary responses may maximise the fitness of individuals in their particular food regimes but will undoubtedly wreak changes to the productivity of whole populations. Some of the responses we observed were not entirely predictable based on existing theory or studies in other systems. Our findings emphasise that evolution will alter and complicate biological responses to global change – with concomitant changes to global food webs that cannot be anticipated based on ecological experiments alone. An important next step is to understand the eco-evolutionary consequences of copepod life history evolution for higher trophic links – how do planktivores that depend on copepods evolve in response to copepod evolution themselves?

## Data accessibility

All data used for analyses and to generate plots are available on request and will be made available on an online public repository prior to publication.

