## Supplementary material for "Copepod life history evolution under high and low food regimes": Figure S2, Figure S3

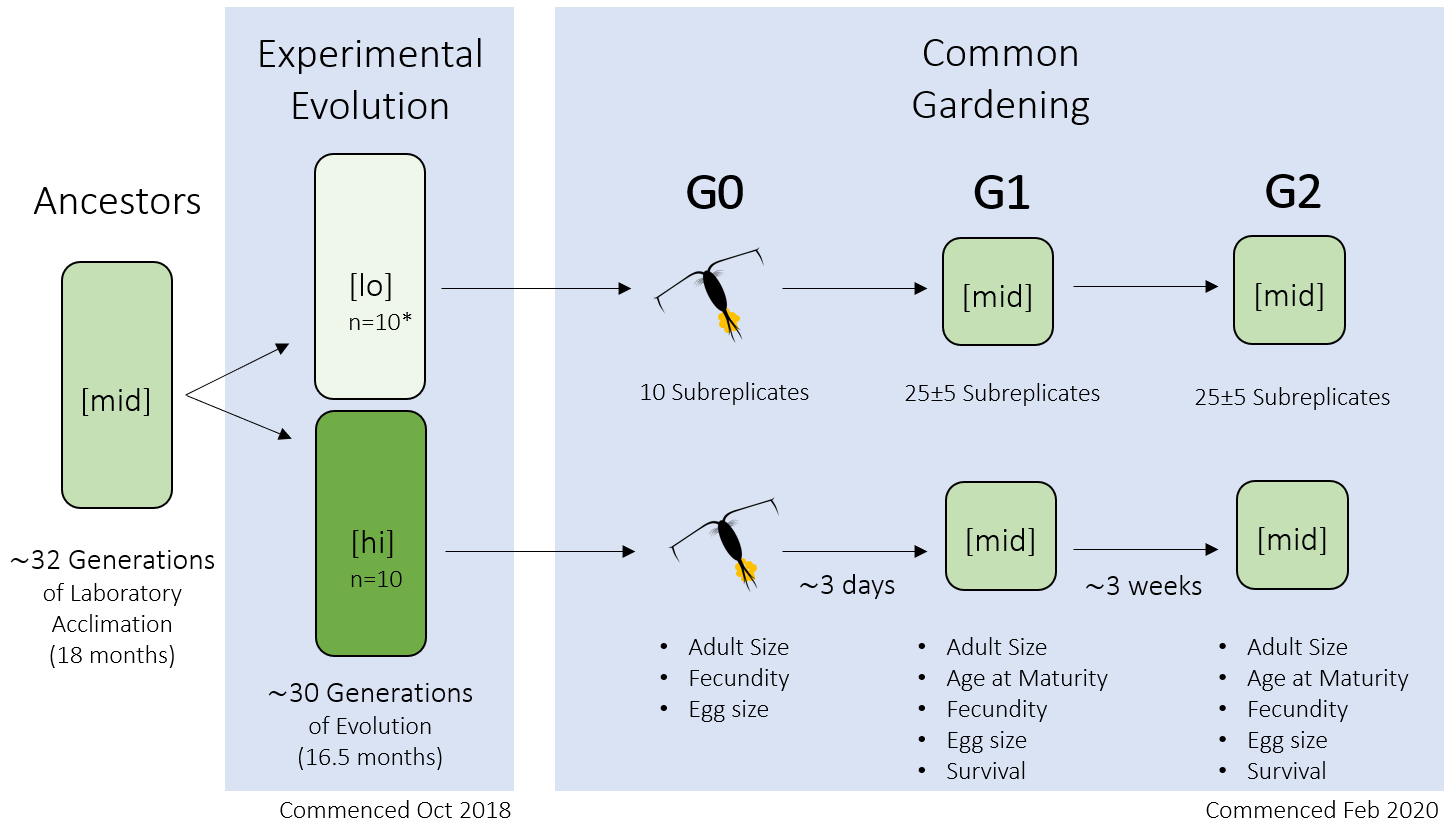

**Figure S2:** Schematic of the series of experiments conducted. Experimental evolution was conducted on 10 populations each of high- and low-food provisioned copepods, for a total of 20 populations. One low-food population went extinct during experimental evolution. Low food (lo) and high food (hi) provisioning different by an order of magnitude, and the intermediate (mid) level of provisioning was the midpoint of high and low. Common gardening began by sampling 10 gravid females (G0) from each population, rearing their G1 offspring to maturity, and in turn rearing the G2 offspring of G1 mothers to maturity. Traits measured in each generation of common gardening are indicated in bullet points. See text for further details.

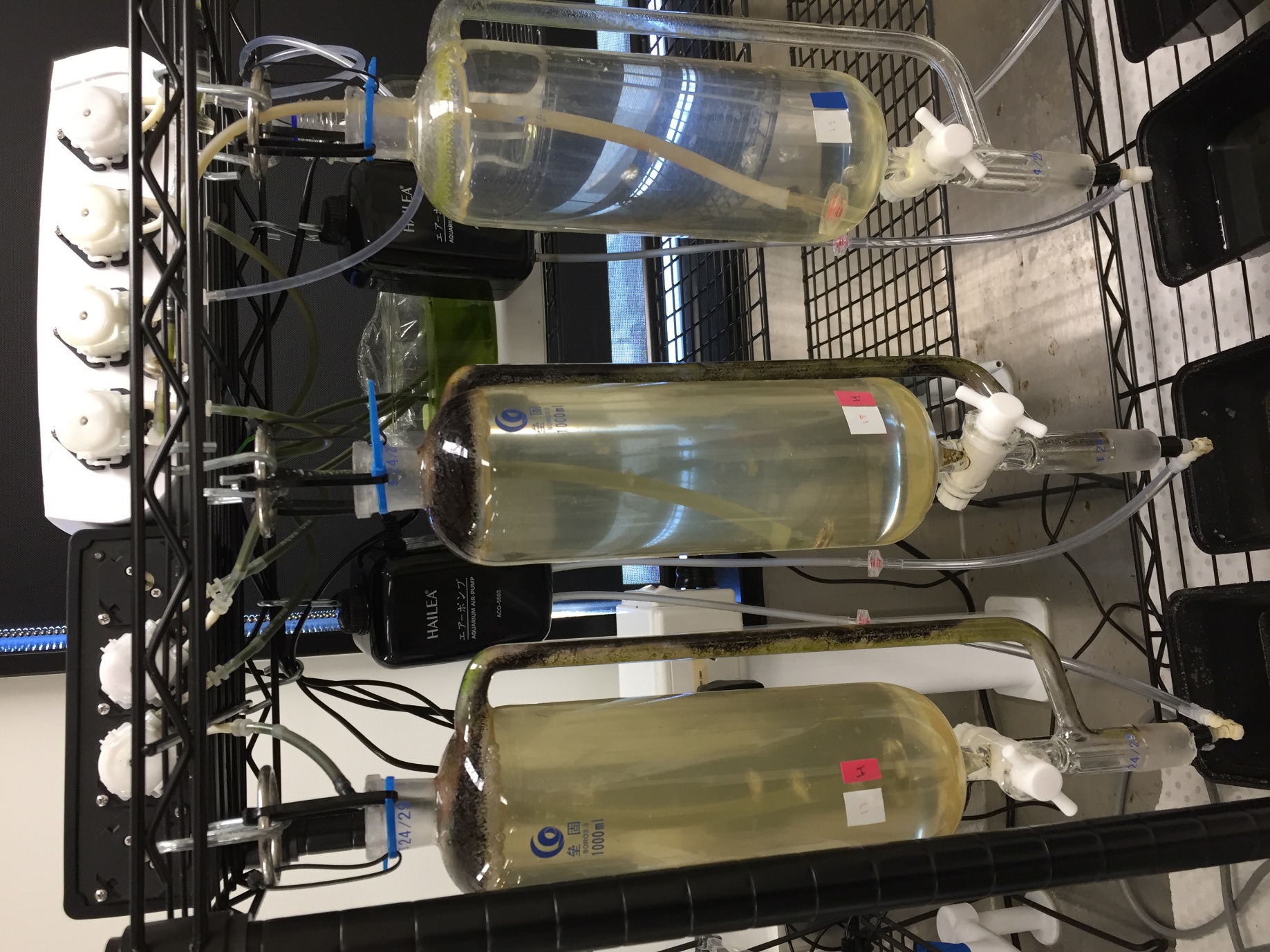

**Figure S3:** 1L glass pressure-equalising dropping funnels used for culturing treatment populations of copepods.
