## Supplementary material for "Copepod life history evolution under high and low food regimes": Figure S1

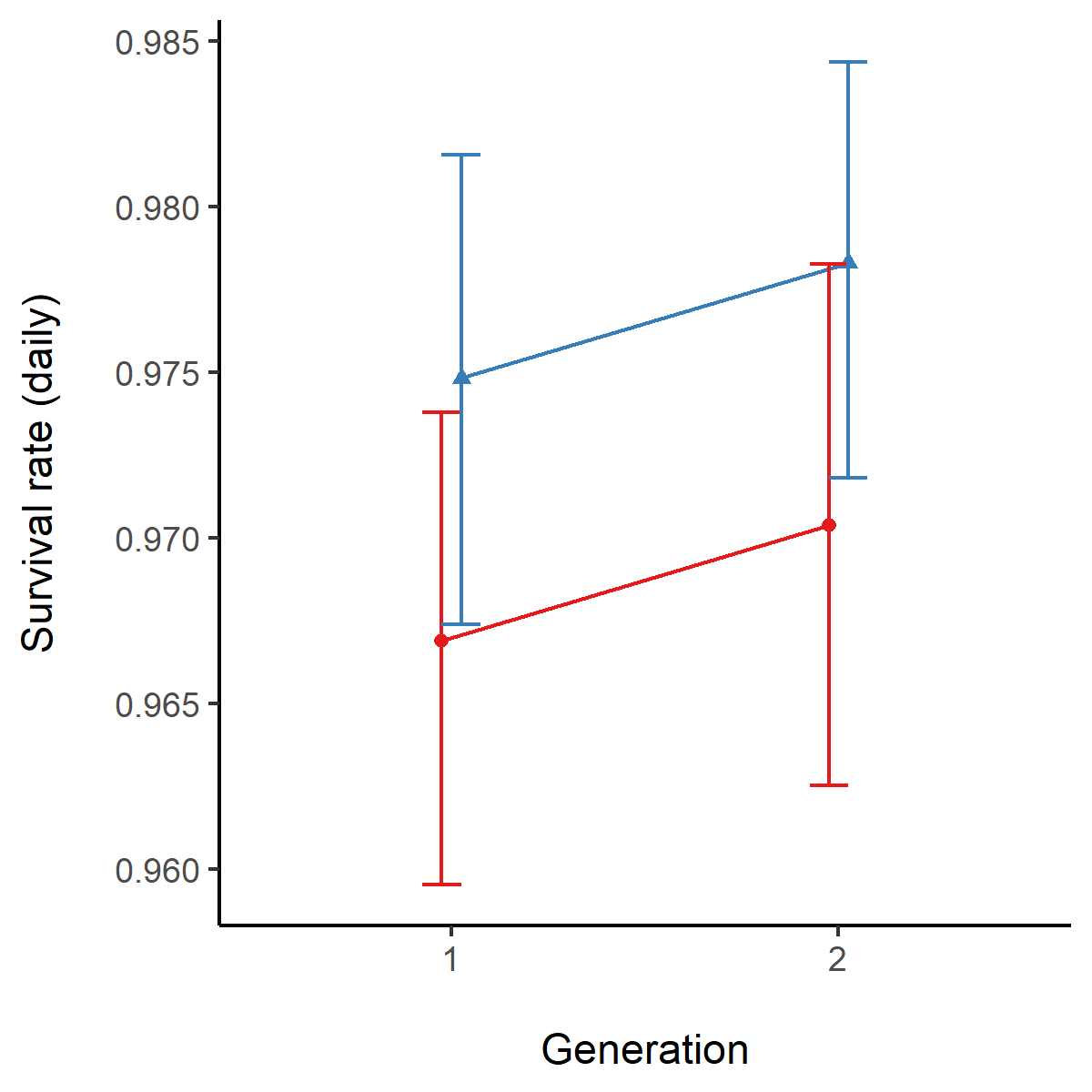

**Figure S1:** Mean daily survival rate, evaluated between hatching to 2 weeks after first observation of sexual maturity (within each replicate) in high-food (solid lines, circular points) and low-food (dotted lines, triangular points) lineages during G1 and G2 of common gardening. Error bars show between-culture bootstrapped 95% confidence intervals.
