## Supplementary material for "Copepod life history evolution under high and low food regimes": Table S1

**Table S1:** Linear mixed effects model for the effect of evolutionary treatment on daily survival rate across two generations (G1, G2), with ancestral G0 populations reared within feeding blocks of four. *p* values are provided for tests of interest, significant effects are specified in bold, and random effect is specified in italics. Degrees of freedom (df) reported as numerator df, denominator df.

| **Effect** | **Df** | **Mean Sq** | **F** | ***p*** |
| --- | --- | --- | --- | --- |
| Treatment | 1, 12.88 | 1.23 x 10^-3^ | 3.53 | 0.08 |
| Generation | 1, 62.04 | 1.96 x 10^-4^ | 0.49 | 0.49 |
| Block | 4, 12.29 | 1.71 x 10^-4^ | 0.54 |  |
| Treat x Gen | 1, 61.34 | 8.75 x 10^-5^ | 0.25 | 0.62 |
| *Culture(Treat)* |  | 3.14 x 10^-4^ |  |  |
